# *ATP7B* Variant c.1934T>G p.Met645Arg Causes Wilson Disease by Promoting Exon 6 Skipping

**DOI:** 10.1101/693572

**Authors:** Daniele Merico, Carl Spickett, Matthew O’Hara, Boyko Kakaradov, Amit G. Deshwar, Phil Fradkin, Shreshth Gandhi, Jiexin Gao, Solomon Grant, Ken Kron, Frank W. Schmitges, Zvi Shalev, Mark Sun, Marta Verby, Matthew Cahill, James J. Dowling, Johan Fransson, Erno Wienholds, Brendan J. Frey

## Abstract

Wilson Disease is a recessive genetic disorder caused by pathogenic loss-of-function variants in the *ATP7B* gene. It is characterized by disrupted copper homeostasis resulting in liver disease and/or neurological abnormalities. The variant NM_000053.3:c.1934T>G (Met645Arg) has been reported as compound heterozygous and is highly prevalent among Wilson Disease patients of Spanish descent. Accordingly, it is classified as pathogenic by leading molecular diagnostic centers. However, functional studies suggest that the amino acid change does not alter protein function, leading one ClinVar submitter to question its pathogenicity. Here we used a minigene system and gene-edited HepG2 cells to demonstrate that c.1934T>G causes approximately 70% skipping of exon 6. Exon 6 skipping results in frameshift and stop gain, which is expected to cause loss of *ATP7B* function. The elucidation of the mechanistic effect for this variant resolves any doubt about its pathogenicity and enables the development of genetic medicines for restoring correct splicing.

## INTRODUCTION

Wilson Disease (OMIM # 277900) is a recessive disorder caused by homozygous or compound heterozygous loss-of-function variants in the *ATP7B* gene (ATPase copper transporting beta; NCBI Gene ID: 540), with an estimated prevalence of 3.3 / 100,000 subjects [Członkowska 2018]. *ATP7B* is a transmembrane copper transporter that primarily exerts its function in liver hepatocytes, where it is required for copper loading onto ceruloplasmin, the main bloodstream copper transporter, and for excess copper excretion into the bile. Insufficient *ATP7B* function results in copper accumulation in hepatocytes, which leads to liver pathology, and also accumulation in other organs such as the brain, which leads to neurological and neuropsychiatric alterations [Członkowska 2018]. Physiologically, copper is bound by extracellular and intracellular chaperones like ceruloplasmin, whereas excess copper often exists in the chaperone-free form. Free copper causes oxidative damage to tissues and leads to Wilson Disease, which is fatal if untreated [Członkowska 2018]. While the standard of care overall improves patient life expectancy and quality of life, 45% of patients exhibit poor long-term adherence because of adverse effects or cumbersome dosing requirements, and 10% of patients with neurological symptoms deteriorate after treatment. Consequently, there is a recognized need for novel and improved therapeutics [Członkowska 2018].

The variant NM_000053.3:c.1934T>G (p.Met645Arg, hg19 genomic coordinates chr13:52535985:A>C) has been reported in several Wilson Disease patients, typically in compound heterozygosity with truncating (i.e. stop-gain, frameshift, large deletions) or missense variants and only once in homozygosity. It was first identified in two Jewish patients in two independent surveys [Shah 1997] [Kalinsky 1998]. It was then identified in 2 / 17 unrelated patients from the Gran Canaria island [Garcia-Villarreal 2000], in 1 / 46 unrelated Brazilian patients [Deguti 2004], in 22 / 40 (55%) unrelated patients from Spain [Margarit 2005], in 1 / 47 unrelated Italian pediatric patients [Nicastro 2009], in 2 unrelated patients of Ecuadorian and Moroccan ethnicity [Lepori 2012] and 1 / 35 patients from Southern Brazil with predominant non-Spanish European descent [Bem 2013]. The Italian pediatric patient was the only homozygous individual [Nicastro 2009]. It is noteworthy that all of the Spanish patients who carried c.1934T>G in compound heterozygosity with another pathogenic variant had hepatic disease; in addition, subjects with a truncating variant in trans had an earlier disease onset (5-14 years old), whereas subjects with a missense variant in trans had more variable onset (7-50 years old) [Margarit 2005]. In contrast, the Brazilian subject was reported to have both liver and brain-related disease, with the latter deemed of greater severity [Deguti 2004]. These details were not available for other reported cases.

Several groups have examined the Met645Arg amino acid substitution and failed to demonstrate a significant effect on protein function. First, copper transport was studied by two different groups using complementary approaches. In Sf9 insect cells infected with a viral vector carrying the *ATP7B* mutant protein cDNA but lacking endogenous copper transporter activity, copper transport was quantified in microsomal vesicles. Copper uptake was normal for the Met645R variant, whereas most of the other pathogenic missense variants that were tested resulted in reduced uptake: 14 / 25 displayed no or low uptake and 10 / 25 displayed partially reduced uptake [Huster 2012]. Second, human SV40-transformed ATP7B-null (YST) fibroblasts were transfected with a plasmid containing the mutated ATP7B-mGFP cDNA, showing that the Met645Arg mutant protein was expressed and was able to transport copper, which is consistent with previous findings [Braiterman 2014]. In addition, vesicular trafficking was characterized by the same group, monitoring migration to the apical membrane (anterograde) in response to elevated copper and return back to the Golgi apparatus (retrograde) in response to depleted copper. Polarized hepatic WIF-B cells were infected with a viral vector carrying the cDNA of the *ATP7B* mutant protein fused with GFP, showing normal anterograde and retrograde trafficking for Met645Arg but not for other pathogenic missense variants [Braiterman 2014]. Last, another group studied the variant effect on *ATP7B* stability mediated by *COMMD1* (Copper metabolism domain containing 1), which binds *ATP7B* and promotes its proteolytic degradation, acting as a negative regulator but also as a quality control mechanism. In HEK293T cells transfected with a plasmid containing the mutated ATP7B-flag cDNA, Met645Arg did not increase *COMMD1* binding. In contrast, increased *COMMD1* binding and reduced *ATP7B* protein expression was observed for other pathogenic missense variants [de Bie 2007]. Based on the studies described above, it is unlikely that the amino acid change is the cause of the observed pathogenicity. Since these studies used cDNA constructs lacking introns they could not investigate effects on splicing.

As of 17 September 2019, 6 / 8 ClinVar submitters have reported the variant as ‘pathogenic’ (Integrated Genetics/Laboratory Corporation of America; Genetic Services Laboratory, University of Chicago; Illumina Clinical Services Laboratory; Fulgent Genetics; Invitae; OMIM) and 1 / 8 as ‘likely pathogenic’ (Counsyl). Only 1 / 8 submitters has classified it as a ‘variant of unknown clinical significance’ (VUS) (SIB, Swiss Institute of Bioinformatics), which was motivated by the negative evidence on the effect of Met645Arg on protein activity (https://www.ncbi.nlm.nih.gov/clinvar/variation/3862/).

We used a minigene system and gene-edited HepG2 cells to demonstrate that c.1934T>G causes approximately 70% skipping of exon 6, as suggested by in-silico analysis. Exon 6 skipping results in frameshift and stop gain, which is expected to cause loss of *ATP7B* function. We therefore propose that c.1934T>G should be classified as a pathogenic or likely pathogenic variant.

## METHODS

### In-silico variant effect analysis

Amino acid impact predictions based on SIFT [SIFT 2001], PolyPhen2-HDIV, PolyPhen2-HVAR [PP2 2010] and Mutation Assessor [MA 2011] were obtained from dbNSFP version v3.5a [dbNSFP 2016].

For splicing, we trained a deep neural network that takes as input a genomic sequence, and predicts the presence of an exon or different types of negatives (e.g. an exon with an incorrect splice site). To construct the training dataset, we used the human reference sequence build 37 and well-supported (TSL = 1) annotated protein-coding exons in Gencode v25 [Frankish 2018] as positive examples. For each positive example, we constructed multiple negative examples that correspond to different missplicing outcomes. The network architecture includes convolutional and locally connected layers, and a recurrent network with long short-term memory along the length of the sequence as the final aggregation layer [Wainberg 2018]. To counteract class imbalance during training, we created mini-batches balanced by the different categories of negatives and trained the classifier using the Adam optimizer [Kingma 2014]. We identified a set of optimal models by running random hyperparameter search and then evaluating performance on a held-out portion of the dataset as well as additional, independent datasets capturing the splicing effect or splicing-mediated pathogenicity of genetic variants [Buratti 2010] [Soemedi 2017] [Landrum 2018]. Finally, we used an ensemble of the top performing models for variant effect prediction.

### Minigene system

Minigene plasmids for *ATP7B* exon 6 were designed to contain part of the *ATP7B* genomic locus, comprising exons 5 to 7 and including the complete sequence of introns 5 and 6. A CMV promoter was placed upstream of exon 5. To minimize aberrant splicing of the transcribed mRNA fragments, consensus splice acceptor sequences of exon 5 were removed by deleting 9 nucleotides from the 5’ end of exon 5; likewise, consensus splice donor sequences of exon 7 were removed by deleting 40 nucleotides from the 3’ end of exon 7. Minigenes were constructed by DNA assembly of PCR fragments that were amplified from HEK293T genomic DNA using KOD Hot Start DNA polymerase (Novagen). For the wild-type minigene, the full fragment was amplified with primer pairs P461 and P463. For the mutant minigene, the c.1934T>G variant was introduced by site-directed mutagenesis of two overlapping fragments which were amplified with primer pairs P461 and P465 and primer pairs P464 and P463 respectively. PCR fragments were cloned into a CMV-containing expression vector, linearized with primers P459 and P460, using a NEBuilder HiFi DNA Assembly Kit (New England Biolabs) according to manufacturer’s instructions. Plasmid DNA was isolated using the Presto Mini Plasmid Kit (Geneaid). Minigenes were used for splicing assays in triplicate in HEK293T, HepG2, HuH-7 and Hep3B cells. Total RNA was isolated 48 hours post transfection using the Qiagen RNeasy kit. For RT-PCR analysis, first strand synthesis was performed using high-capacity cDNA kit with 500ng of RNA and random primers. PCR was performed with primers P243 and P472 and separated on 2% agarose gels stained with 0.05% Redsafe (FroggaBio). See Supplementary Dataset 1 for primer sequences.

### Cell culture, maintenance and transfection

HEK293T, HepG2 and Hep3B cell lines were purchased from ATCC. Huh-7 cells were purchased from JCRB. HEK293T cells were maintained in IMDM (Gibco) supplemented with 10% COSMIC serum (ThermoFisher), L-glutamine (2 mM) (Gibco) and Penicillin/Streptomycin (100 units/ml) (Gibco). Hep3B and HepG2 cells were maintained in DMEM 4.5 g/L glucose (10% FBS (Gibco) and Penicillin/Streptomycin). Huh-7 cells were maintained in DMEM 1 g/L glucose (10% FBS and Penicillin/Streptomycin). Cells were dissociated by incubation with TrypLE (ThermoFisher) and then equal parts media was added to the suspension. The cells were pelleted by centrifugation at 200 x g for 5 minutes and the supernatant was removed. Cells were then resuspended and diluted for correct plating densities. All cells were reverse transfected with 500 ng minigene plasmid DNA with Lipofectamine 3000 and P3000 reagent (Invitrogen) in 12 well plates. 4.5 x 10^5^ HEK293T cells were seeded per well. 3 x 10^5^ cells per well were seeded for HepG2, Hep3B and Huh-7.

### Gene-editing of HepG2 cells and screening

The 2F3 edited HepG2 clone, compound heterozygous for c.1934T>G and a loss-of-function re-arrangement, was created by the Centre for Commercialization of Regenerative Medicine (CCRM) (Toronto, Canada) using the CRISPR/Cas9 editing system and a pEGFP-puro plasmid for co-selection of transformants (see Supplementary Figure 3). Single-cell derived clones were screened for the presence of the c.1944T>G variant by digital droplet PCR followed by Sanger sequencing. A positive clone was selected and the presence of the c.1944T>G variant was re-confirmed by Sanger sequencing at The Centre for Applied Genomics (TCAG) (Toronto, Canada). For this, genomic DNA was extracted using gSYNC DNA Extraction kit (Geneaid) and quantified by nanovue (GE Healthcare). A touchdown PCR was performed on 200 ng of extracted genomic DNA with primers P488 and P650 to amplify a 759 bp fragment of *ATP7B* containing exon 6 and surrounding intronic regions. PCR was checked using UCSC in silico PCR. PCR products were purified using the PCR cleanup kit (Geneaid) following the manufacturer’s instructions. DNA was submitted for Sanger sequencing at TCAG with forward and reverse PCR primers. Finally, the clone was selected and submitted for whole genome sequencing (WGS).

The 2A1 clone, homozygous for a T insertion at the cut site resulting in complete loss-of-function, was also generated by CCRM and genotyped following the same methodology, except for the final WGS step.

The 1F6 and 1E8 clones (homozygous for c.1934T>G) were created using the CRISPR/Cas9 editing system at Deep Genomics. One million parental HepG2 cells were nucleofected with crRNAs/tracrRNA complex, ssODN and Cas9 endonuclease (IDT) using the Lonza Nucleofector 4D platform according to the manufacturer’s recommended protocol (cell line solution SF and program EH-100). Oxford Nanopore MinION sequencing was performed to verify editing efficiency. Cells were then diluted and plated at dilution in 96 well plates to isolate individual clones. Individual clones were screened by Sanger sequencing to identify edited populations. These were further confirmed by sequencing of amplicons on the MiSeq platform. See Supplementary Dataset 2 for crRNA and template sequences.

### Whole genome sequencing and analysis of the 2F3 HepG2 clone

PCR-free whole genome sequencing (WGS) libraries were prepared at The Center for Applied Genomics (TCAG) with Illumina’s Nextera DNA Flex Library Prep Kit, using genomic DNA from wild-type and c.1934T>G edited HepG2 cells. Both WGS libraries were sequenced to 40x mean coverage on Illumina HiSeq X. WGS alignment and variant calling was performed using BWA-MEM v0.7.12 [Li 2010] and a custom version of the human reference genome (b37) extended with an extra chromosome containing the sequence of the pEGFP-puro plasmid used for co-selection of transformants. Variant calling was performed using GATKv3.7 [DePristo 2011]. Reads aligned to the plasmid or within 2kb of its integration breakpoints on chr13 were collected, adapter- and quality-trimmed using Trim_Galore v.0.4.4 (https://www.bioinformatics.babraham.ac.uk/projects/trim_galore/), and de-novo assembled using SPAdes v3.9.0 [Bushmanova 2018] with k-mer sizes of 21, 33, 55, 77, 99. As independent validation of the plasmid integration, gDNA primers were designed to PCR amplify and Sanger sequence fragments spanning the putative breakpoints and a 20 nt region with no WGS coverage in the middle of the plasmid sequence. See Supplementary Dataset 1 for primer sequences.

### qPCR

12 replicates of wild-type, 2F3, 1F6 and 1E8 edited HepG2 cells were seeded into 12-well plates (3.5 x 10^5^ cells per well). Total RNA was extracted from the cells using the RNeasy mini-kit with Qiacube automated extraction (Qiagen). DNase was performed on-column with RNase-free DNase kit (Qiagen). RNA concentration and quality were determined by Bioanalyzer (Agilent). All RNA samples had RNA integrity numbers (RIN) above 9. First strand synthesis was performed with 1 µg of RNA using the High Capacity cDNA kit with random primers (Applied Biosystems). For qPCR, primers were designed to amplify *ATP7B* transcripts containing exons 5, 6 and 7 by spanning exon junctions (P2065, P2066). TBP primers were used for normalisation (P378, P379). qPCR was performed with Quantstudio 5 instrument (Thermofisher) with SYBR green reagents (Thermofisher). Cycle thresholds were determined by QuantStudio software, which were used to calculate relative quantity (RQ) of *ATP7B* transcripts by relative standard curve method. See Supplementary Dataset 1 for primer sequences.

### Western blot

Total protein was isolated from wild-type, 2A1, 2F3, 1E8 and 1F6 edited HepG2 cells. Cells were first rinsed with 1 x DPBS (Gibco) and lysed with ice cold RIPA Buffer (Thermofisher) containing HALT protease inhibitors (Thermofisher). Cell lysate was chilled on ice for 10 minutes and centrifuged at 16000 x g to pellet and remove cell debris. Protein amounts were determined by BCA assay (Pierce). Samples were heated at 70°C with NuPAGE sample buffer containing NuPAGE sample reducing agent (Thermofisher). 20 µg of total protein was separated on a 10% TRIS-BIS (NuPAGE) gel at 200V for 1 Hour. Separated proteins were transferred to PVDF membrane at 350 mA constant for 90 minutes using the Mini-TransBlot system (Bio-Rad). Transfer was validated by ponceau stain (Sigma). Membranes were blocked in 5% Milk TBST for one hour at room temperature. PVDF membrane was cut at 75 kDa according to protein ladder (BlueElf, FroggaBio). The higher molecular weight portion of the membrane was incubated with *ATP7B* antibody (Abcam, Ab131208) (1/500 in 5% Milk TBST) and the lower molecular weight portion of the membrane was incubated with beta-actin antibody as a loading control (Abcam Ab8227) (1/1000 in 5% Milk TBST) overnight at 4°C. Samples were washed three times for 10 minutes in TBST and then incubated in secondary antibody (70745 anti-rabbit HRP conjugated, 1/5000 in 5% Milk TBST) (Cell Signalling) at room temperature for 1 hour. The membranes were washed three times for 10 minutes in TBST. The images were taken after a 30 second exposure for beta-actin and after a 1 minute exposure for *ATP7B* using the Amersham Imager 680 using Amersham ECL reagents (GE).

## RESULTS

### Overview

We carried out a thorough in-silico analysis of the c.1934T>G variant effect, which suggested that the amino acid substitution Met645Arg can be tolerated but the variant causes > 50% exon 6 skipping, resulting in frameshift and stop gain. The in-silico splicing prediction was backed by experimental high-throughput data and was further validated in a mini gene system. In addition, we used CRISPR/Cas9 to obtain one clone of HepG2 cells compound heterozygous for c.1934T>G and a large insertion, and two clones of HepG2 cells homozygous for c.1934T>G; qPCR transcript quantification showed 15% of correctly-spliced wild-type transcript level in the compound het and 31-33% in the homozygous clones. Evaluation of c.1934T>G frequency suggests that it is consistent with pathogenicity. We therefore propose that c.1934T>G should be classified as a pathogenic or likely pathogenic variant.

### In-silico analysis of c.1934T>G effect

First, we validated that NM_000053 is the most appropriate principal transcript for *ATP7B* in hepatocytes. NCBI Homo sapiens RefSeq annotation release 109 has five curated isoforms: NM_000053 (21 exons, encoding 1465 amino acids), NM_001243182 (22 exons, encoding 1354 amino acids), NM_001330578 (20 exons, encoding 1387 amino acids), NM_001330579 (19 exons, encoding 1381 amino acids), NM_001005918 (17 exons, encoding 1258 amino acids). In the ClinVar data downloaded in January 2019, NM_000053 is used for 209 / 210 submissions of pathogenic or likely pathogenic variants. Accordingly, based on annotation, protein domain composition and conservation evidence, NM_000053 is categorized as principal by APPRIS 2019_02.v29 [Rodriguez 2013]. In contrast, NM_001243182, NM_001330578 and NM_001330579 are categorized as alternative, whereas NM_001005918 is categorized as minor. Manual review of junctional counts from GTEx v7 [GTEx 2013] reveals that ENST00000242839 comprises the best supported junctions (see Supplementary Figure 1). ENST00000242839 is almost identical to NM_000053, since they differ by only a few nucleotides at the 3’UTR exon 21 end. In conclusion, multiple lines of evidence support using NM_000053 as the principal transcript for *ATP7B* in hepatocytes. Therefore, we used NM_000053.3 as reference transcript throughout the paper.

The Met645 amino acid position is located immediately before a predicted transmembrane element according to UNIPROT (https://www.uniprot.org/uniprot/P35670), which is consistent with the overall protein domain diagram of *ATP7B* showing exon 6 encoding a flexible linker between the copper binding domains and the first transmembrane element [Członkowska 2018]. According to the multiple genome alignments provided by the UCSC hg19 genome browser (https://genome.ucsc.edu/), Met645 is frequently substituted by valine, leucine, alanine and threonine in mammalians and by lysine, glutamine or aspartic acid in other vertebrates; armadillo is the only mammalian showing substitution by arginine (see Supplementary Figure 2). Consistent with this, in-silico amino acid effect predictors return impact scores for Met645Arg that correspond to lack of deleteriousness: 0.239 for SIFT, 0.146 for PolyPhen2-HDIV, 0.119 for PolyPhen2-HVAR, -1.22 for Mutation Assessor [SIFT 2001] [PP2 2010] [MA 2011]. In conclusion, in-silico evidence suggests that this amino acid substitution can be tolerated, or perhaps produce a modest loss of function, in line with results from functional experiments [de Bie 2007] [Huster 2012] [Braiterman 2014].

Next, we assessed if c.1934T>G could alter splicing. c.1934T>G is outside of the splice site consensus region (13 bp from the donor site), but it could still impact exon recognition by altering a splicing silencer or enhancer [Scotti 2016]. We used a deep neural network predictor trained to recognize exons from genomic data, and applied it to score the reference and mutant sequence to obtain a variant score. The predictor is based on a deep learning convolutional neural network that automatically learns to identify sequence motifs important for splicing. This resulted in a predicted change of exon strength corresponding to > 50% exon 6 skipping. In addition, exon skipping quantification based on the high-throughput MaPSy minigene assay was available for this variant: exon 6 inclusion was decreased to 35.5% wild-type in the in-vivo data-set and 53.8% wild-type in the in-vitro data-set [Soemedi 2017].

Since exon 6 is out-of-frame, its skipping leads to a shifted reading frame and the introduction of a stop codon in exon 8 (TGA at c.2259, genomic coordinate chr13:52532543-52532541), which is expected to cause reduced expression via nonsense-mediated decay (NMD). It is particularly compelling that both in-silico and experimental high-throughput data suggest a splicing alteration. This prompted us to further validate this finding experimentally and to accurately quantify the amount of residual correctly-spliced transcript.

### c.1934T>G causes exon skipping in a minigene system

We constructed both a wild-type and variant-containing minigene with exon 6 flanked by full introns and partial exons 5 and 7. Splicing assays were performed in triplicate in four different cell lines by transient minigene transfection. RT-PCR showed partial exon skipping in the wild-type (c.1934T) minigene in all four cell lines (see Figure 1), which could be caused by less-efficient splicing of minigene RNA. Importantly, however, c.1934T>G minigene had almost complete (100%) exon skipping in all cell lines and always exceeded the wild-type by a good margin, thus confirming the c.1934T>G effect on exon skipping.

**Figure 1:**
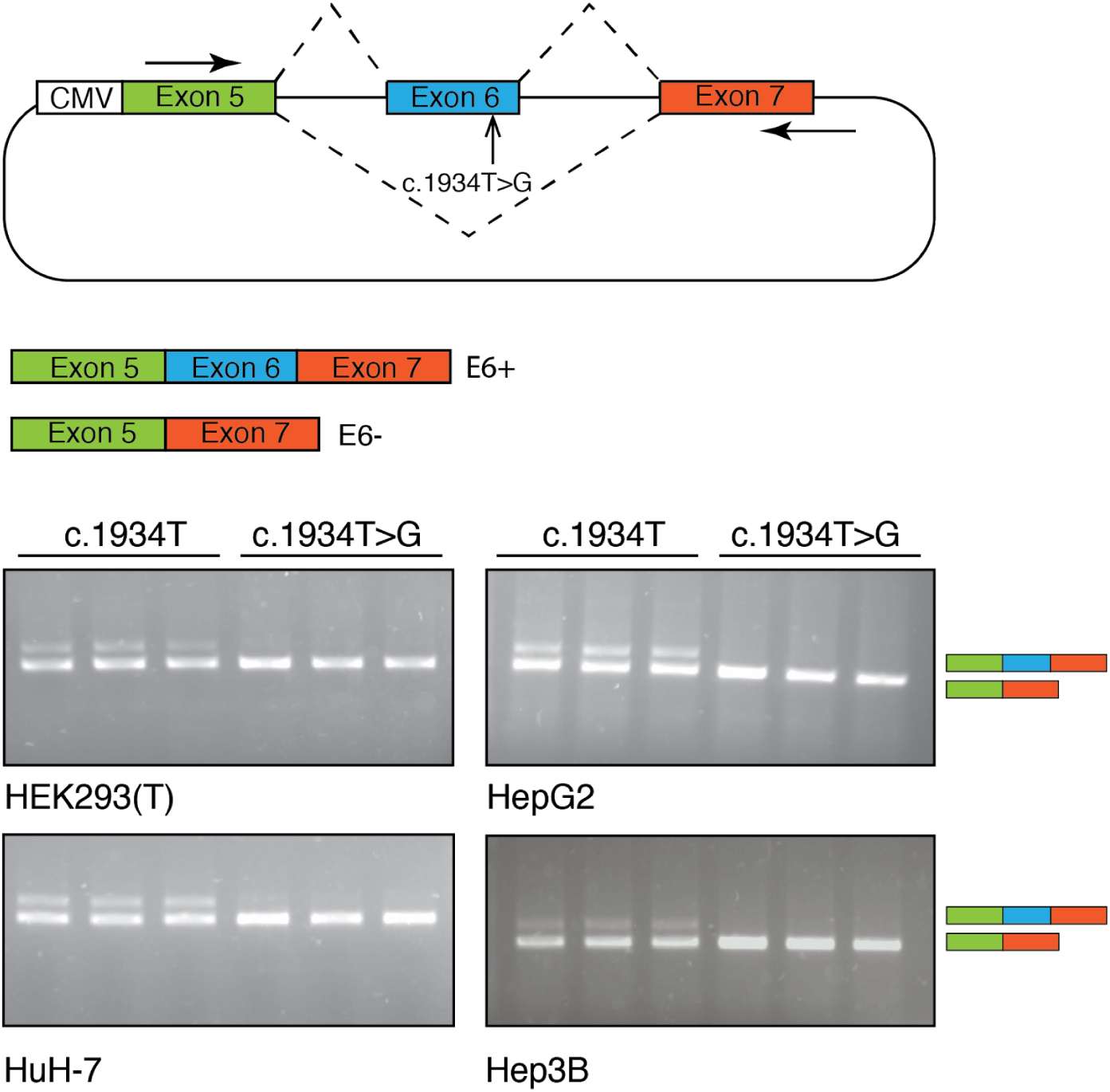
minigene analysis of c.1934T>G variant in four different cell lines (HEK293T, HepG2, HuH-7 and Hep3B). The wild-type minigene (c.1934T) showed partial exon 6 inclusion, whereas the mutant minigene (c.1934T>G) showed almost complete skipping in all four cell lines.

### Generation of c.1934T>G edited HepG2 cells

We used CRISPR/Cas9 to obtain HepG2 cell lines with a c.1934T>G allele to study the variant effect in the endogenous gene (see Table 1). Digital droplet PCR followed by Sanger sequencing were used to screen for clones positive for the c.1934T>G variant.

**Table 1:**
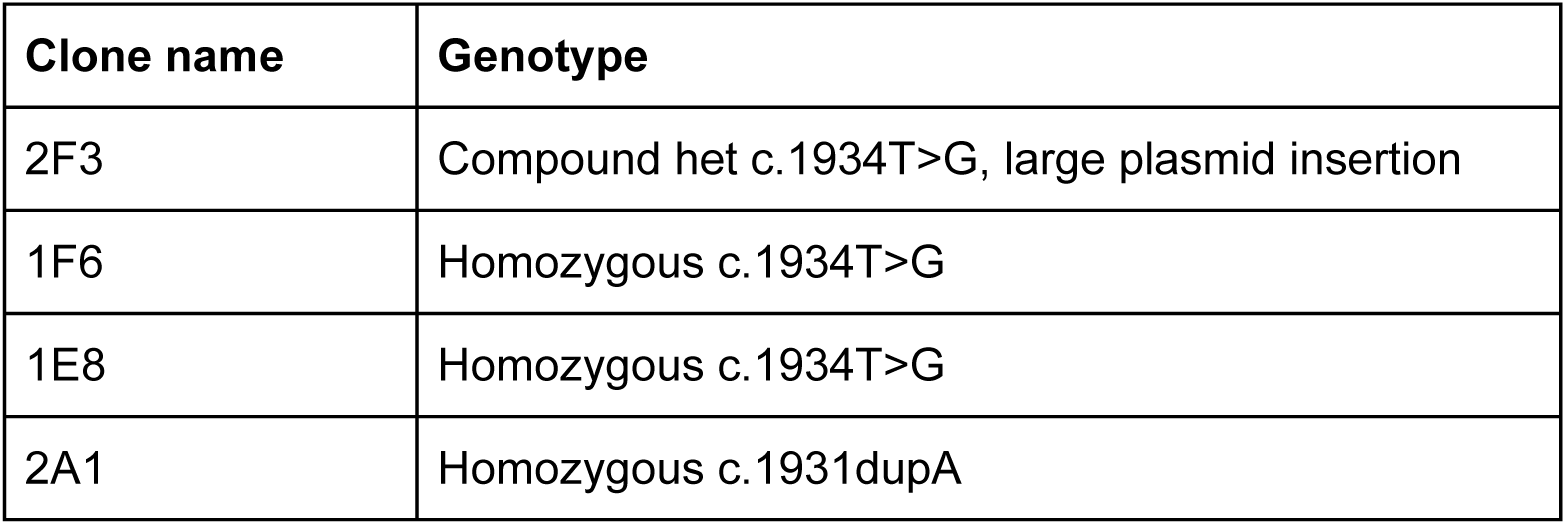
edited HepG2 cells used in this study.

| Clone name | Genotype |
| --- | --- |
| 2F3 | Compound het c.1934T>G, large plasmid insertion |
| 1F6 | Homozygous c.1934T>G |
| 1E8 | Homozygous c.1934T>G |
| 2A1 | Homozygous c.1931dupA |

We first generated one clone compound heterozygous for c.1934T>G. The genotype of this compound heterozygous clone (2F3) was further characterized by whole genome sequencing (WGS). WGS was supplemented with targeted genomic PCR to carefully reconstruct the two alleles at the *ATP7B* locus. WGS alignment to the human reference sequence revealed three major read clusters at the *ATP7B* exon 6 locus. One corresponded to the c.1934T>G allele, showing no other edits or alterations. The other two clusters, corresponding to the second allele, had one portion of the reads aligning to the reference sequence, whereas the other portion of the reads was completely different than the human reference sequence and instead corresponded to a section of the co-transfection plasmid (see Supplementary Figure 3; see Supplementary Figure 4 for the plasmid structure). Based on these results, we inferred the presence of a partial exon 6 duplication flanking a plasmid sequence insertion; we used genomic PCR to validate the breakpoints joining human reference sequence to plasmid sequence (see Supplementary Figure 3; see Supplementary Figure 5 for genomic PCR results). To resolve the full length of the plasmid insertion, we aligned all reads to the human reference extended by the plasmid sequence as an additional chromosome, and retrieved all reads mapping to the plasmid and to 2 kb of reference sequence around the breakpoints; we then performed de-novo assembly and identified two contigs, which included the human-plasmid breakpoints but were separated by a small assembly gap when compared to the original plasmid sequence (see Supplementary Figure 3). We hypothesized that the small assembly gap was due to a region with low sequencing coverage, and we confirmed this by genomic PCR spanning the two contig ends (see Supplementary Figure 5 for genomic PCR results). Therefore, we concluded that this clone presents one allele with a clean c.1934T>G edit, whereas the other allele has a partial duplication of exon 6 and a ∼ 5 kb plasmid insertion.

A second round of editing generated two clones homozygous for c.1934T>G variant (1E8 and 1F6).

### c.1934T>G causes exon skipping and reduced protein expression in edited HepG2 cells

Since exon 6 is out-of-frame, transcripts lacking exon 6 may be degraded by NMD. We thus reasoned that qPCR, with primers overlapping the exon 5-6 and exon 6-7 junctions, would be the most suitable approach to quantify the c.1934T>G splicing effect. In addition, we monitored the qPCR product length to exclude the presence of aberrant transcripts produced from the plasmid insertion allele in 2F3 cells and not degraded by NMD, which are expected to have a different length. RNA was isolated from 12 individual preparations of HepG2 wild-type, 2F3, 1F6 and 1E8 cells. First-strand synthesis and qPCR were used to calculate the relative quantity of transcripts containing exons 5, 6 and 7 between wild-type and edited cells. qPCR results showed that, compared to wild-type cells, compound heterozygous HepG2 cells (2F3 clone) express only 15% of transcript containing exons 5, 6 and 7 (see Figure 2) and homozygous cells (clones 1E8 and 1F6) express only 31-33%. Melt curves for 2F3 cells were indicative of a single PCR product without detection of aberrant transcripts derived from the plasmid insertion (See Supplementary Figure 6).

**Figure 2:**
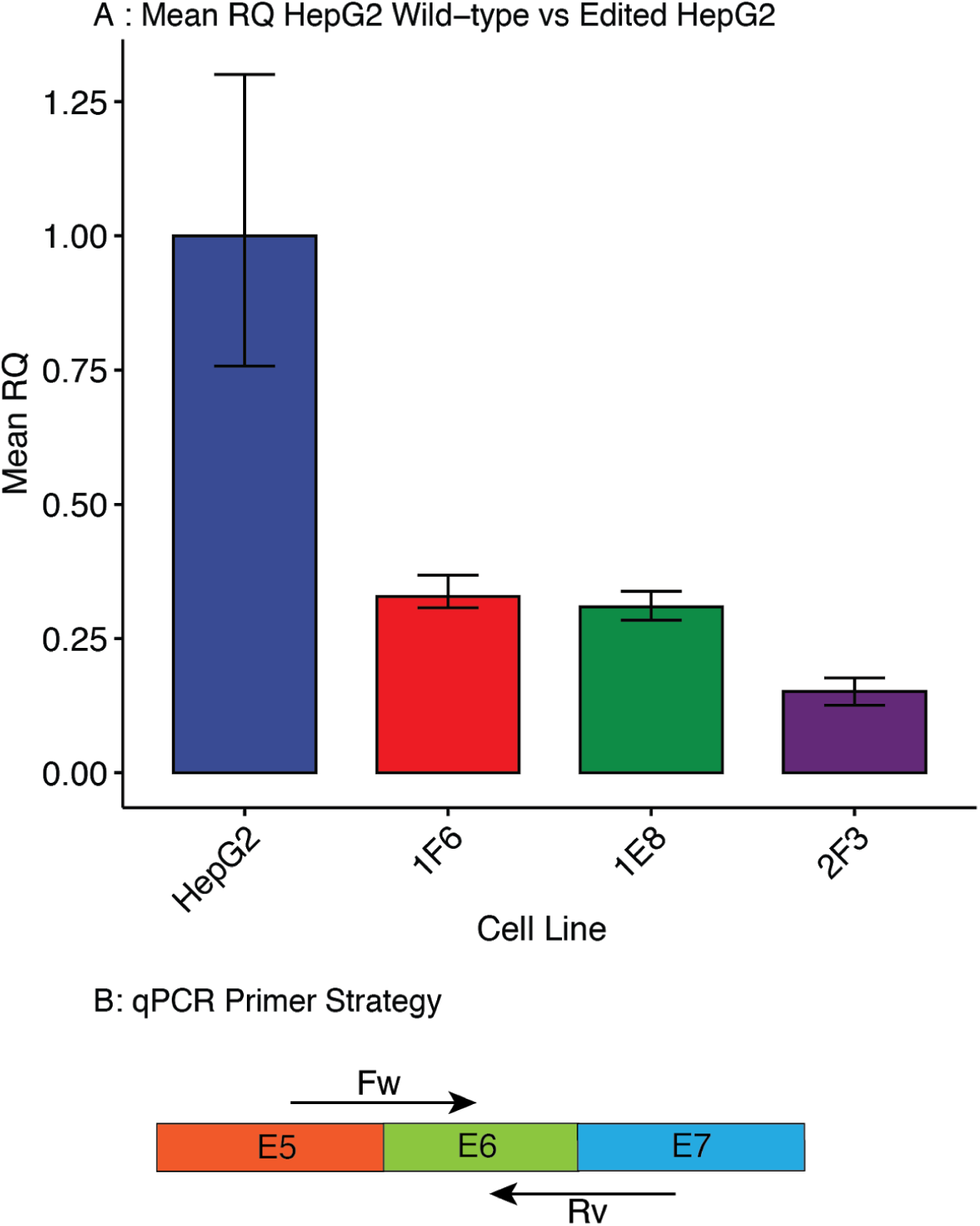
qPCR for relative quantity (RQ) of transcripts containing exons 5, 6 and 7 between wild-type HepG2, 2F3, 1F6 and 1E8 cell lines. (A) Compared to wild-type cells, 2F3 cells have only 15% of exon 5, 6, 7 spanning transcript. 1E8 and 1F6 cells have 31% and 33% respectively compared to wild-type cells. The barplot displays the mean RQ of 12 independent RNA extractions for each cell line, with error bars corresponding to minimum or maximum RQ. (B) PCR strategy with forward (FW) and reverse (RV) primers spanning the boundaries between exons 5 and 6 and 6 and 7 respectively.

In addition, protein lysates were extracted from wild-type and edited HepG2 cells to determine *ATP7B* protein expression by western blot. Compared to wild-type cells, compound heterozygous (2F3) and homozygous (1E8, 1F6) cells express reduced levels of ATP7B (see Figure 3), with increased *ATP7B* expression in 1E8 and 1F6 homozygous cell lines compared to 2F3 cells. A control 2A1 *ATP7B* knock-out cell line has no detectable levels of ATP7B.

**Figure 3:**
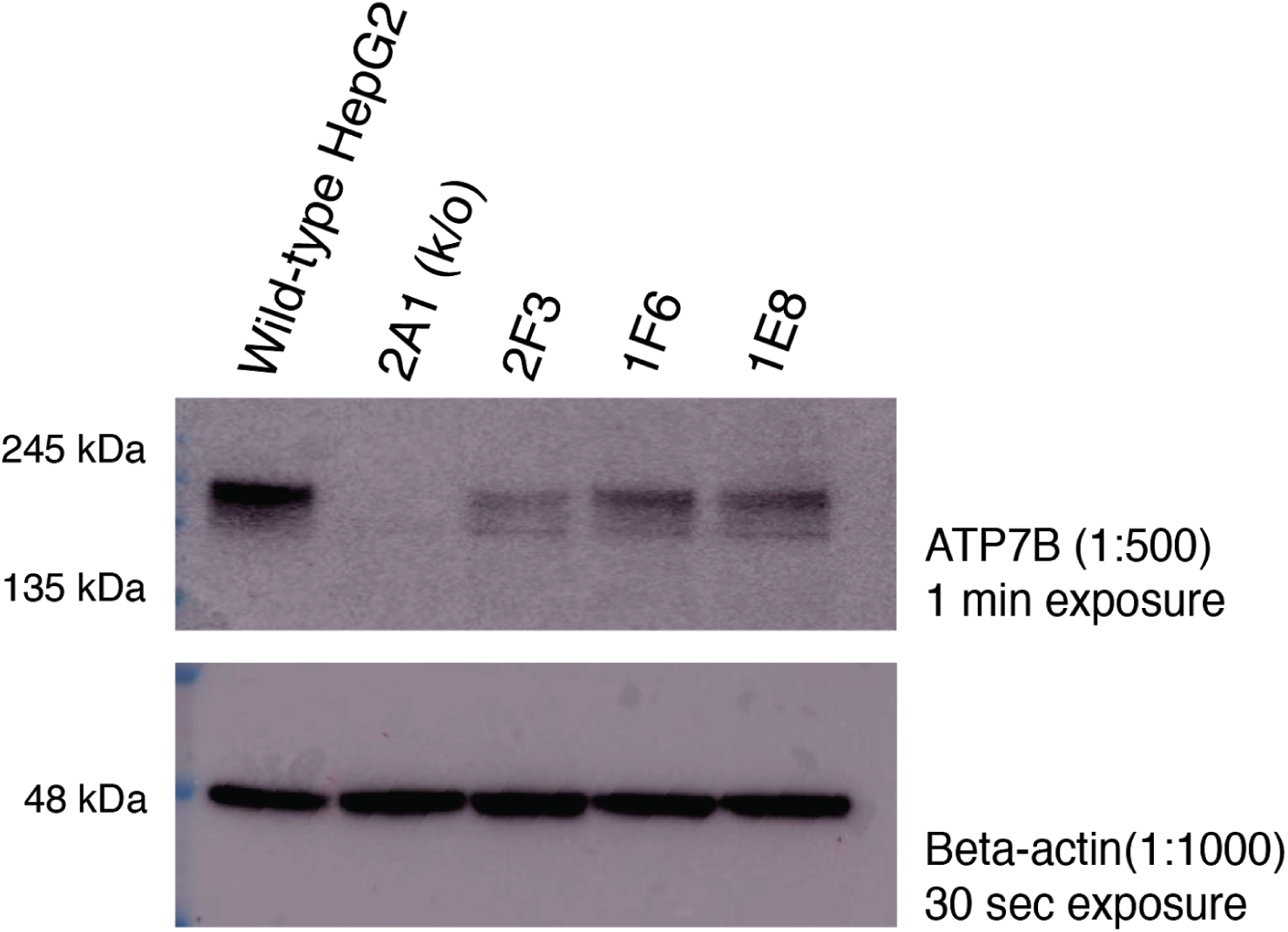
Western blot performed on total lysate(s) from wild-type and edited HepG2 cells. Immunoblot for *ATP7B* shows decreased protein expression in 2F3, 1F6 and 1E8 cell lines compared to HepG2 wild-type cells. Increased expression in 1F6 and 1E8 homozygous lines compared to 2F3 compound heterozygous line (null allele by large insertion). No observed expression in the 2A1 knock-out (k/o) cell line. Immunoblot for beta-actin was used as a protein loading control.

### c.1934T>G frequency is consistent with pathogenicity

A large survey of Wilson Disease in Spain reported c.1934T>G in 55% of the patients [Margarit 2005], but did not report the variant frequency in the general Spanish population. The best matched gnomAD populations appears to be Latino, with frequency = 0.0022330, followed by Southern European, with frequency = 0.0004342. A direct comparison of the allele frequency in this disease cohort (allele count = 22 / 80) to the gnomAD allele frequency in the Latino population (allele count = 79 / 35376) results in a Fisher’s Exact Test p-value = 2.05 x 10^−38^ and OR = 169 (95% confidence interval = 94-296).

However, it may be over-optimistic to use these allele frequencies for a direct case-control comparison because the population ethnicity is only partially matched. Assuming that c.1934T>G is typically pathogenic only in compound heterozygosity with other pathogenic variants, and that Wilson Disease prevalence in Spain is the same as the consensus prevalence of 3.3 / 100,000, it is possible to estimate the c.1934T>G allele frequency in the general population of Spain, and verify if it is consistent with expectations, by solving the following equations.

#### Definitions

- Frequency of c.1934T>G: *Fv* (to be estimated)
- Total frequency of *ATP7B* pathogenic variants: *Ftot* (to be estimated)
- Population of Spain: *Spain_pop* = 46.7 x 10^6^
- Population of Spain with Wilson Disease: *Spain_WD_pop* = *Spain_pop * WD_prevalence*
- Prevalence of Wilson Disease: *WD_prevalence =* 3.3 x 10^−5^
- Fraction of Spanish WD patients with c.1934T>G: *Spain_WD_var* = 0.55

#### Equations

- 2 * (Fv * (Ftot - Fv)) * Spain_pop = Spain_WD_pop * Spain_WD_var
- (Ftot ^ 2 - Fv ^ 2) * Spain_pop = Spain_WD_pop
- Spain_WD_pop = WD_prevalence * Spain_pop

Solving these equations results in the Spanish c.1934T>G frequency *Fv* = 0.002355 and total pathogenic variant frequency *Ftot* = 0.006209. Strikingly, this Spanish c.1934T>G frequency estimate is very similar to the c.1934T>G Latino frequency in gnomAD (0.002233).

It is worth considering that the fraction of patients with c.1934T>G is subject to stochastic sampling error; we can repeat our calculation to estimate the frequency of c.1934T>G using instead the Clopper-Pearson 70% confidence interval (0.4559, 0.6412) of *Spain_WD_var*, which leads to Freq (c.1934T>G) = 0.001775-0.003075. Compared to the gnomAD Latino frequency, these values appear to be reasonably in line with expectations.

In conclusion, c.1934T>G is present at a much higher frequency in Wilson Disease patients of Spanish descent and its allele frequency in the general population of Spanish descent is consistent with the consensus prevalence of Wilson Disease.

### Proposed updated ACMG classification

The experimental evidence that c.1934T>G reduces the intact transcript to about 30% normal levels supports the ACMG [Richards 2015] pathogenicity evidence *PS3*, i.e. functional studies supportive of damaging effect on the gene or gene product. The frequency of c.1934T>G in Wilson Disease patients compared to the general population supports the ACMG pathogenicity evidence *PS4*, i.e. prevalence in affected individuals significantly increased compared to controls, which could be downgraded to *PM* to account for residual uncertainties (see the previous section for details). *PS3* and *PS4* result in the classification *pathogenic* based on the rule *≥ 2 Strong (PS1-PS4)*, whereas if *PS4* is replaced by *PM* the resulting classification is *likely pathogenic* based on the rule *1 Strong (PS1-PS4) and 1-2 Moderate*.

## DISCUSSION

NM_000053.3:c.1934T>G (p.Met645Arg) has been reported in 55% of Spanish patients with Wilson Disease and in several patients of Jewish or Southern European descent. It has clearly higher frequency among individuals with Wilson Disease compared to the general population. The mutant Met645Arg protein has been studied in several functional assays, none of which revealed significant differences compared to the wild-type protein. Accordingly, in-silico analysis suggests that this amino acid substitution could be tolerated. This has raised some concerns about the pathogenicity of this variant.

In-silico analysis and previously published high-throughput data suggest that NM_000053.3:c.1934T>G causes > 50% exon 6 skipping. Since exon 6 is out-of-frame, skipping is expected to result in loss of *ATP7B* function. Minigene constructs confirmed c.1934T>G causes exon 6 skipping in four different cell lines. In addition, edited HepG2 cells compound heterozygous for c.1934T>G and a large plasmid insertion present a major reduction of *ATP7B* expression: qPCR detected 15% correctly spliced transcript and Western blot showed almost complete loss of protein expression. Accordingly, qPCR detected 31-33% correctly spliced transcript in edited HepG2 cells homozygous for c.1934T>G. These results suggest that c.1934T>G causes Wilson Disease chiefly by altering splicing and reducing *ATP7B* expression.

The paucity of c.1934T>G homozygotes among reported Wilson Disease cases may be explained by the relatively high correctly spliced transcript in homozygotes (about 30%). It is thus possible that c.1934T>G homozygous patients have later-onset liver disease (e.g. 40-50 years old) and are less likely to be diagnosed for Wilson Disease. However, the only one reported homozygous case belonged to a pediatric group with median age at diagnosis of 7.4 years and maximum age at diagnosis of 21 years [Nicastro 2009]. It is also possible that homozygotes require additional risk factors (genetic or not) to develop Wilson Disease, such as common hypofunctional variants impacting *ATP7B*. Anyway, these results suggest that c.1934T>G homozygotes are very unlikely to present a more severe or earlier onset disease than compound heterozygotes with a truncating variant.

In addition to dispelling doubts on c.1934T>G pathogenicity, the elucidation of the splicing effect enables the development of genetic medicines for splicing modulation. Steric blocking antisense oligonucleotides (SBOs) have been successfully used for splicing restoration in-vitro as well as in-vivo [Havens 2016]. The chemical formulation with a phosphorothioate backbone and 2’OMe or 2’MOE modifications has proven particularly safe and effective, with, as of 2019, one FDA-approved drug (nusinersen, for spinal muscular atrophy) [Finkel 2017], one approved investigational new drug (QR-110 for Leber congenital amaurosis) [Dulla 2018] and one successful tailor-made clinical application [Yu 2018]. Highly effective hepatocyte uptake can be mediated by GalNAc cluster conjugation [Dowdy 2017]. The standard of care for Wilson Disease presents serious adverse effects and adherence issues [Członkowska 2018], thus c.1934T>G homozygous and compound heterozygous patients could benefit from a novel SBO therapeutic restoring *ATP7B* exon 6 inclusion.

## ACKNOWLEDGEMENTS

We would like to thank the staff of the Centre for Commercialization of Regenerative Medicine (CCRM, Toronto) for the HepG2 CRISPR/Cas9 editing, and the staff of The Centre for Applied Genomics (TCAG, Toronto) for the edited HepG2 whole genome sequencing and bioinformatics analysis. We would also like to thank Dr. Rachel Soemedi and Prof. William G. Fairbrother for providing access to MaPSy predictions. Finally, we would like to thank Dr. Frederick K. Askari for fruitful discussions about Wilson Disease.

## SUPPLEMENTARY ITEMS

### List of supplementary figures

**Supplementary figure 1:**
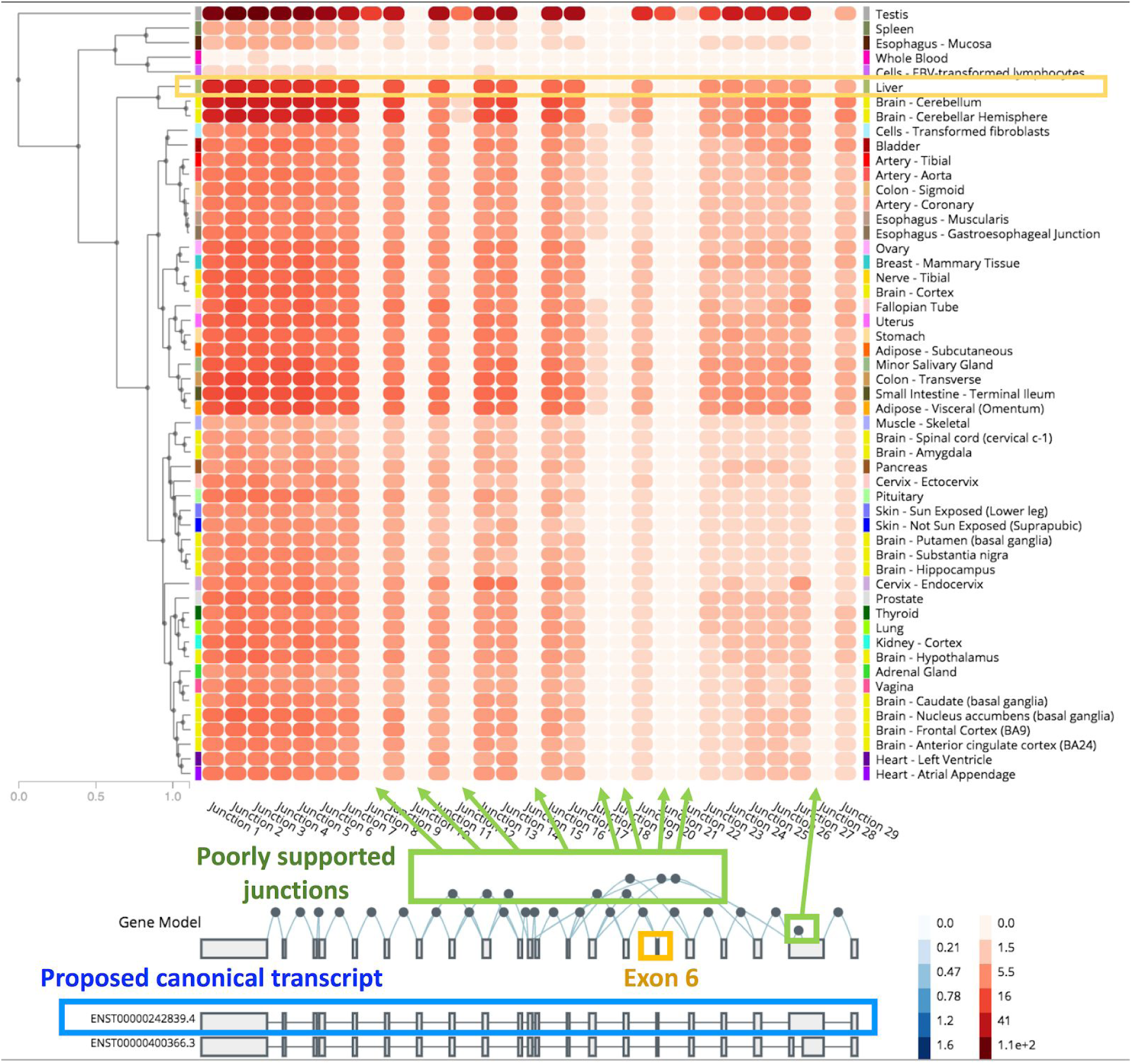
manual analysis of GTEx junctions suggests ENST00000242839 is the highest expressed isoform in liver.

**Supplementary figure 2:**
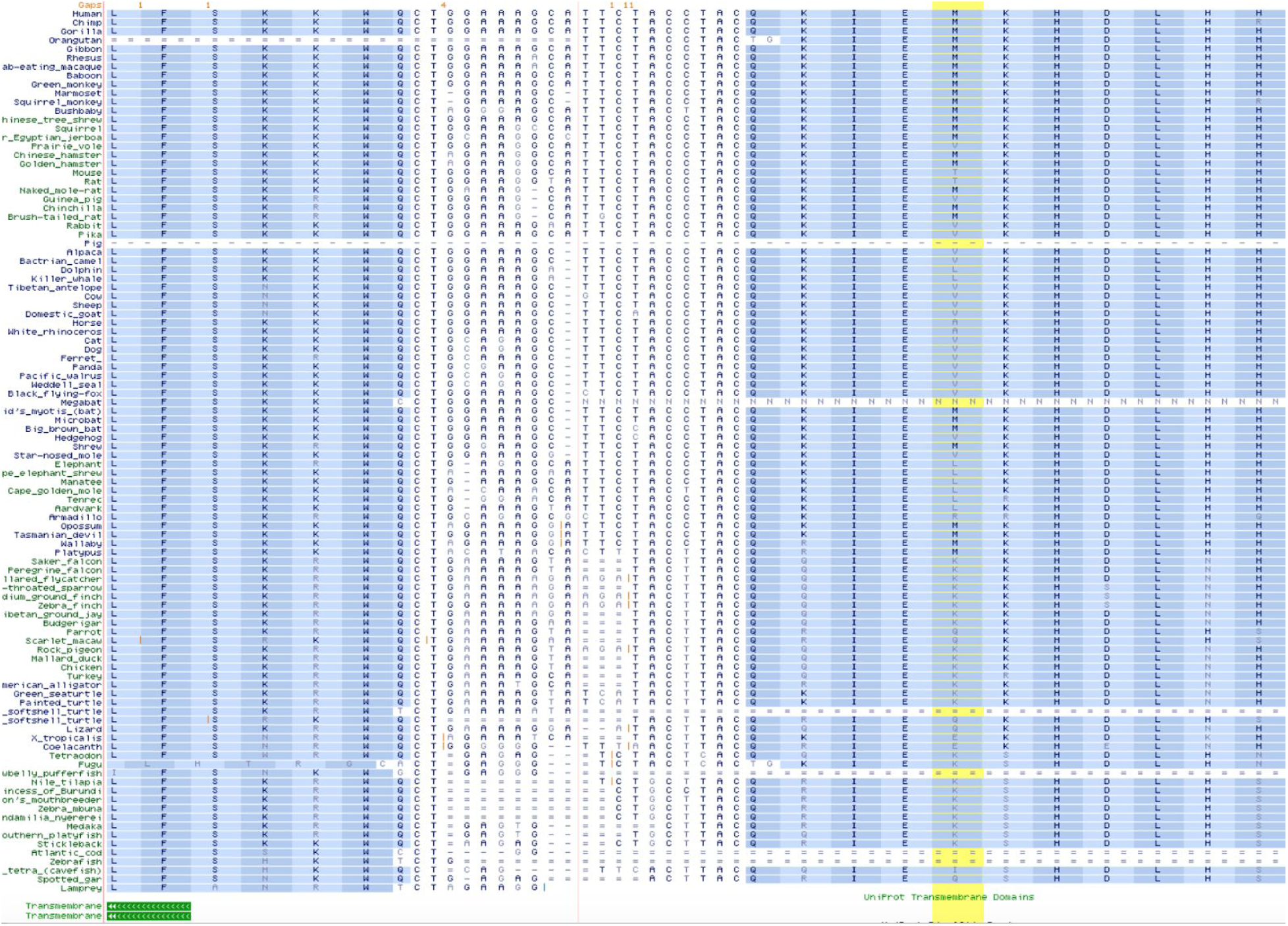
UCSC browser 100 vertebrate multiple sequence alignment showing amino acid substitution patterns for Met645 (exon 6 is on the left, exon 7 is on the right).

**Supplementary figure 3:**
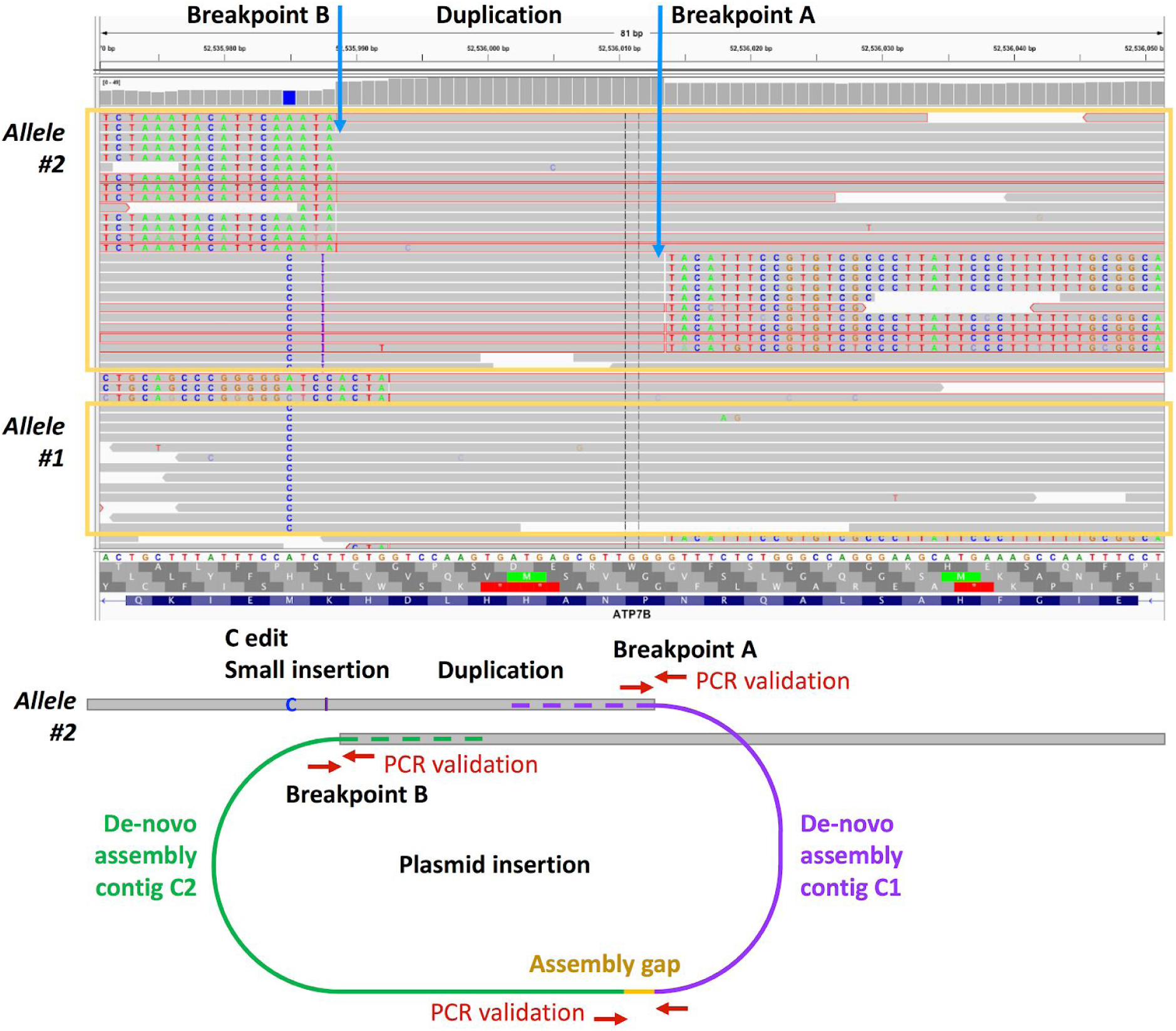
WGS results for the 2F3 HepG2 clone. Top, alignment to the human reference sequence (visualized using the Integrative Genomics Viewer [IGV 2017]) reveals three major read clusters, one corresponding to the edited c.1934T>G allele (allele 1) and the others suggesting a partial exon 6 duplication and plasmid insertion (allele 2). Bottom, the reconstructed sequence of allele 2, showing the human genome reference sequence as gray blocks, the contigs obtained by de-novo assembly as green and violet lines, the assembly gap as a gold line, the PCR primer sets used for validation as red arrow pairs (see Supplementary Figure 5 for PCR results); note that the plasmid insertion length is not proportional to the exon 6 length and that the de-novo assembly contigs span the exon 6 as well as nearby genomic reference sequence as suggested by the dashed lines.

**Supplementary figure 4:**
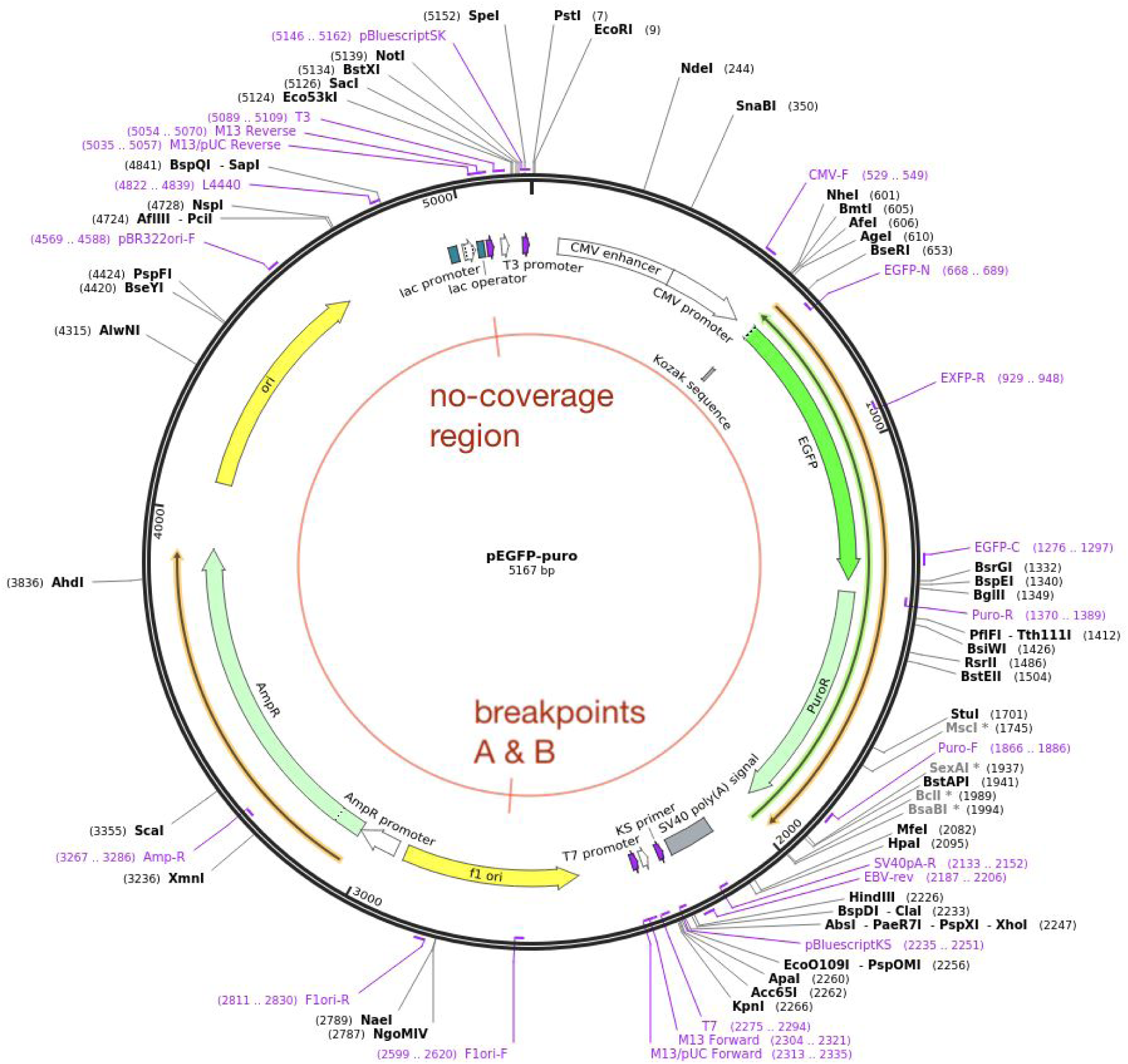
pEGFP-puro plasmid diagram, indicating the position of breakpoints A and B, and the “no coverage region” leading to the assembly gap after de-novo assembly.

**Supplementary figure 5:**
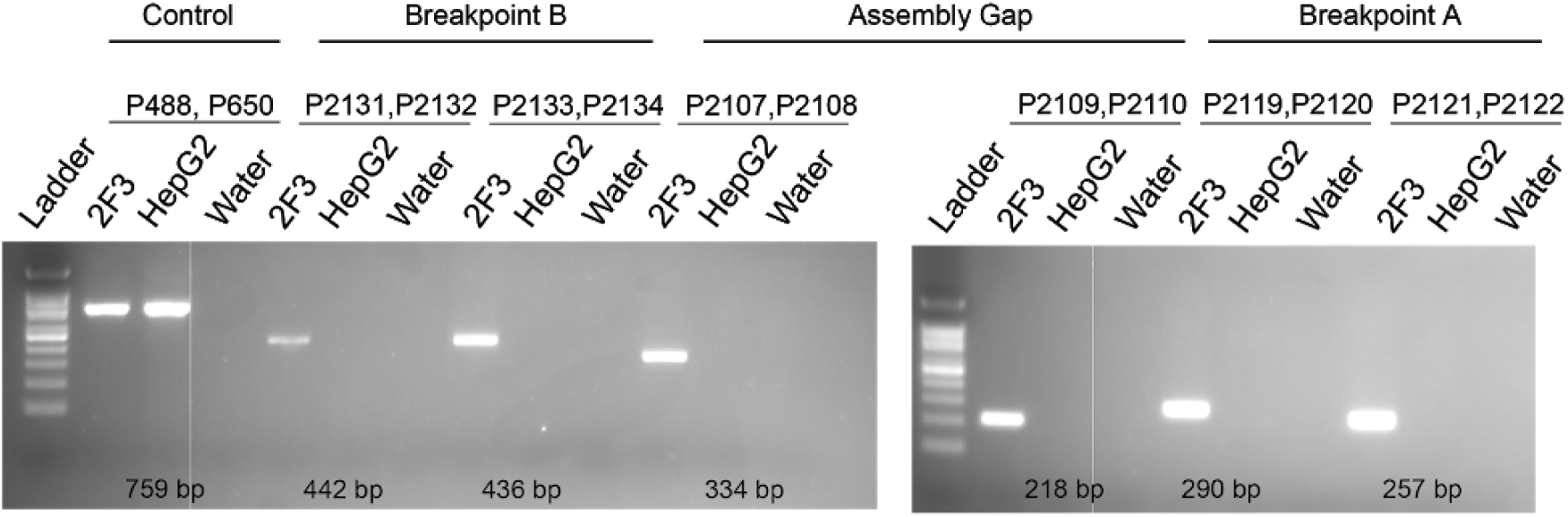
genomic PCR covering Breakpoints A, B and assembly gap according to Figure 2 show amplification specifically in edited cells. Control PCR for ATP7B amplified in both edited and wild-type cells. Water serves as a negative template control.

**Supplementary Figure 6:**
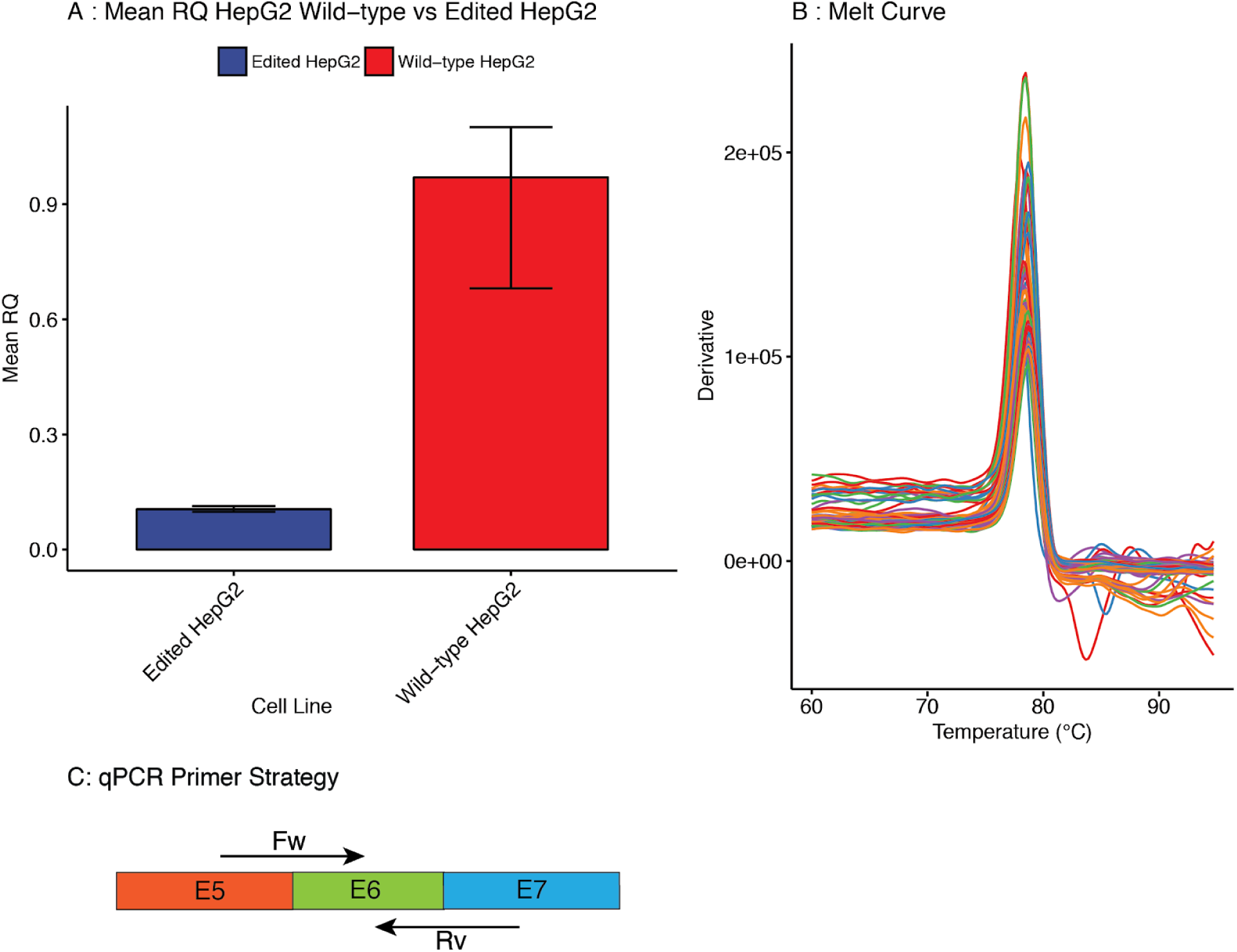
qPCR for relative quantity (RQ) of transcripts containing exons 5, 6 and 7 between wild-type HepG2 and edited 2F3 cells. (A) Compared to wild-type cells, edited HepG2 cells have reduced exon 5, 6, 7 spanning transcript; the barplot displays the mean RQ of 12 independent RNA extractions for each cell line, with error bars corresponding to minimum or maximum RQ. (B) Melt curve for all biological replicates is indicative of a single PCR product. (C) PCR strategy with forward (FW) and reverse (RV) primers spanning the boundaries between exons 5 and 6 and 6 and 7 respectively.

### List of supplementary datasets

- Supplementary Dataset 1: primers
- Supplementary Dataset 2: guide RNA and templates for 1F6 and 1E8 editing

